# Mapping of centriolar proteins onto the post-embryonic lineage of *C. elegans*

**DOI:** 10.1101/2023.05.25.542230

**Authors:** Nils Kalbfuss, Antonin Berger, Pierre Gönczy

**Affiliations:** Swiss Institute for Experimental Cancer Research (ISREC), School of Life Sciences, Swiss Federal Institute of Technology Lausanne (EPFL), Lausanne, Switzerland, CH-1015

**Keywords:** *C. elegans*, centriole, peri-centriolar material (PCM), cell fusion, lineage

## Abstract

Centrioles, together with the surrounding peri-centriolar material (PCM), constitute the centrosome, a major microtubule-organizing center of animal cells. Despite being critical in many cells for signaling, motility and division, centrioles can be eliminated in some systems, including in the vast majority of differentiating cells during embryogenesis in *Caenorhabditis elegans*. Whether the cells retaining centrioles in the resulting L1 larvae do so because they lack an activity that eliminates centrioles in the other cells is not known. Moreover, the extent to which centrioles and PCM remain present in later stages of worm development, when all cells but those of the germ line are terminally differentiated, is not known. Here, by fusing cells that lack centrioles with cells that retain them, we established that L1 larvae do not possess a diffusible elimination activity sufficient to remove centrioles. Moreover, analyzing PCM core proteins in L1 larval cells that retain centrioles, we found that some such proteins, but not all, are present as well. Furthermore, we uncovered that foci of centriolar proteins remain present in specific terminally differentiated cells of adult hermaphrodites and males, in particular in the somatic gonad. Correlating the time at which cells were born with the fate of their centrioles revealed that it is not cell age, but instead cell fate, that determines whether and when centrioles are eliminated. Overall, our work maps the localization of centriolar and PCM core proteins in the post-embryonic *C. elegans* lineage, thereby providing an essential blueprint for uncovering mechanisms modulating their presence and function.

## Introduction

The centrosome is a major microtubule-organizing center (MTOC) of animal cells, comprising centrioles and surrounding pericentriolar material (PCM), which is enriched in the γ-tubulin ring complex and thereby nucleates microtubules (Moritz et al., 1995). Centrosomes are important for various aspects of cellular architecture and function, including polarity and faithful segregation of chromosomes at cell division (reviewed by Marshall, 2007).

PCM components can redistribute to other subcellular locations in some instances. This is the case in vertebrate myotubes, where PCM components surround the nuclear periphery, thereby forming non-centrosomal MTOCs (Bugnard et al., 2005; Tassin et al., 1985). Similarly, in *Caenorhabditis elegans* embryos, PCM components such as the γ-tubulin ring complex components GIP-1 and MZT-1, as well as MTOC activity, localize at the apical surface of polarized intestinal cells (Feldman & Priess, 2012; Sallee et al., 2018). Moreover, whereas centrioles are required to initiate sensory cilium formation in *C. elegans*, they disappear during axoneme assembly, leaving merely PCM components in an “acentriolar centrosome” at the ciliary base (Garbrecht et al., 2021; Magescas et al., 2021; Serwas et al., 2017).

Centrioles are eliminated in other contexts, including during oogenesis of metazoan organisms, where such elimination ensures that the resulting zygote is endowed with only the two centrioles contributed by the sperm. Moreover, we found recently that during *C. elegans* embryogenesis, centrioles are eliminated in a stereotyped manner in ∼88 % of cells, leaving only 68 cells harboring one or two foci containing the centriolar proteins SAS-4::GFP and GFP::SAS-7 in the L1 larval stage that follows (Kalbfuss and Gönczy, 2023). Whether such centriolar foci also retain PCM components is not known, nor it is known whether centrioles are present later in worm development in terminally differentiated cells.

Centrioles in *C. elegans* are minute cylindrical organelles with a 9-fold radial symmetry of peripheral microtubules, the molecular architecture of which has been resolved (Woglar et al., 2022) (Fig. 1A). Of particular relevance for this study, the evolutionarily conserved centriolar protein SAS-4 localizes inside the microtubule wall, whereas SAS-7 localizes more peripherally, as does SPD-2, which is also a PCM core component. Likewise, the PCM core proteins PCMD-1 and SPD-5, which surround centrioles throughout the cell cycle as does SPD-2 (Erpf et al., 2019; Hamill et al., 2002; Kemp et al., 2004), are present outside the microtubule wall. The PCM expands as cells approach mitosis, when it becomes enriched in further proteins, including the γ-tubulin ring complex components GIP-1 and MZT-1 (Hannak et al., 2002; Sallee et al., 2018; Strome et al., 2001). During mitosis in the early embryo, additional proteins, including the Polo-like kinase PLK-1, are recruited to the centrosome, further enhancing microtubule nucleation (Chase et al., 2000; Ohta et al., 2020).

**Figure 1:**
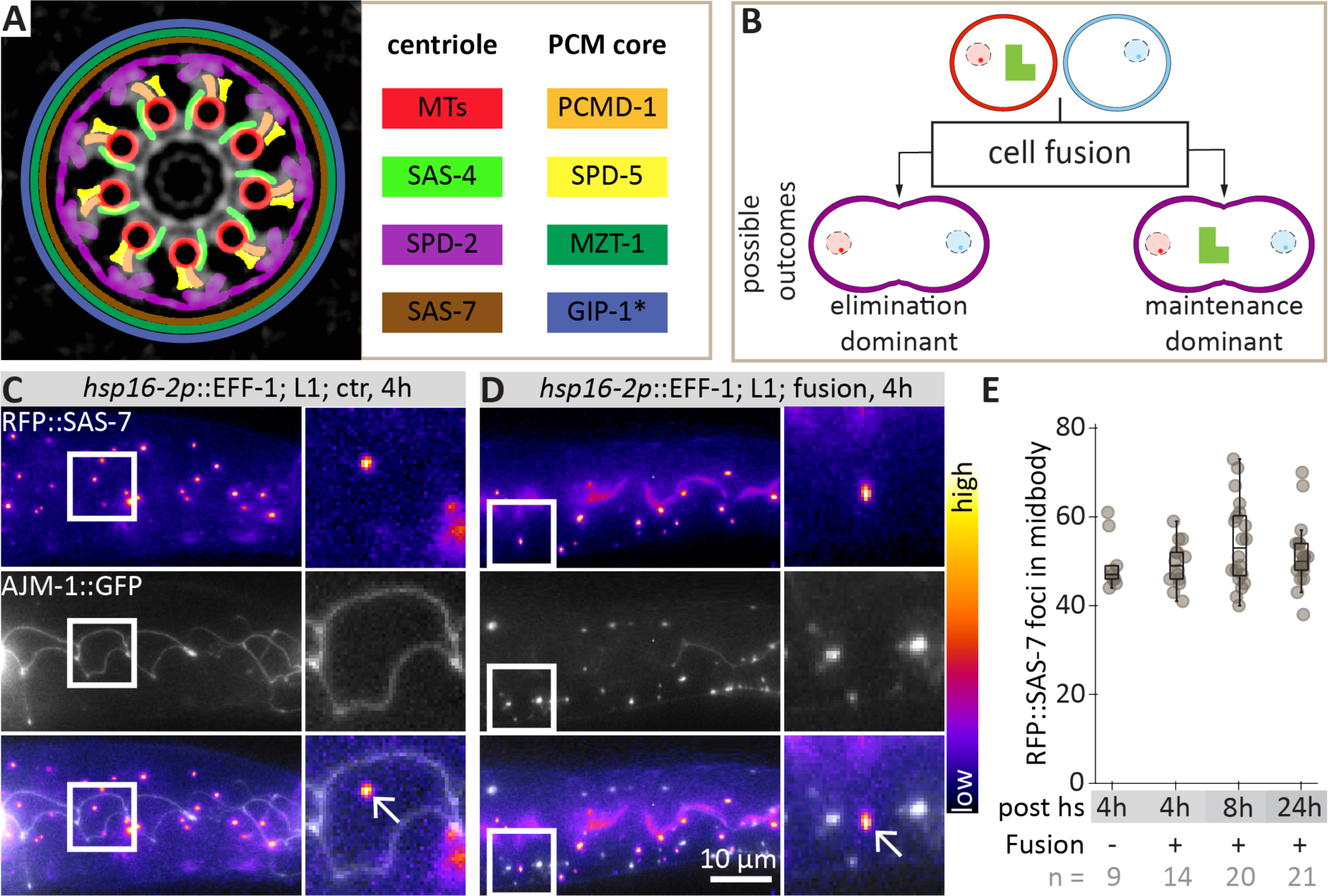
Fusion experiments reveal that centriole elimination is not constantly active in L1. **(A)** Schematic of centriole and PCM core components (modified from Woglar et al., 2022). The asterisk for GIP-1 indicates approximate localization as indicated by the distribution of the GIP-1-interacting protein TBG-1 (Woglar et al., 2022), which is compatible with the actual approximate localization of GIP-1 (Sallee et al., 2018). **(B)** Possible outcomes of fusion between a cell that maintains centrioles (red) and a cell that does not (blue). If centriole elimination is dominant, centriole number should decrease after fusion, whereas it should remain the same if centriole maintenance is dominant. Note that in principle fusion could also lead to *de novo* centriole formation – a process that has not been reported in *C. elegans* and is therefore omitted from the schematic for simplicity. **(C, D)** Maximum z-projection wide-field microscopy of live paralyzed L1 larva expressing RFP::SAS-7 and the cell-cell junction marker AJM-1::GFP outlining seam cells, either without heat-shock (C) or 4 hours after heat-shock (D). Only planes containing seam cells are shown. Boxed regions are shown as magnified insets on the right. Arrows point to RFP::SAS-7 foci of seam cells. In D, note disappearance of cell-cell junctions marked by AJM-1::GFP, indicating successful fusion, but maintenance of RFP::SAS-7 foci (arrows). **(E)** Number of RFP::SAS-7 positive cells in midbody (i.e. between intestinal-pharyngeal and intestinal-rectal valve) 4, 8 and 24 hours after heat-shock. The experiments at 4 hours were conducted with *hsp16-2p*::EFF-1 RFP::SAS-7 AJM-1::GFP, later time points with *hsp16-2p*::AFF-1 RFP::SAS-7 AJM-1::GFP *scmp*::GFP. Pairwise comparisons with the non-heat-shocked control using Wilcoxon-Mann-Whitney test after Bonferroni correction were all non-significant (p-value = 1).

Seven of the 68 cells that retain centrioles in L1 larvae, the rectal epithelial cells B, F, U, Y, K’, as well as the amphid socket cells AmsoL/R, have terminally exited the cell cycle but nevertheless retain a focus of SAS-4::GFP and GFP::SAS-7 (hereafter referred to as “non-proliferating” cells) (Kalbfuss and Gönczy, 2023). The remaining 61 cells are blast cells that will later proliferate. Whether further elimination of centrioles occurs later in development when the progeny of these proliferating cells has terminally exited the cell cycle in turn has not been addressed.

In this study, we first show through experimental cell fusion that L1 larvae do not possess a diffusible centriole elimination activity that is sufficient to remove centrioles. Furthermore, we report that centrosomes in the non-proliferating cells that retain centrioles in L1 larvae also retain some PCM core proteins, but not all. Thereafter, we analyze the presence of centrioles in terminally differentiated somatic cells of the adult hermaphrodite and male, finding that centrioles are maintained in some somatic gonadal cells, in a manner that does not depend on cell age, but instead on cell type.

## Material and Methods

### Strains

*C. elegans* strains were cultured according to standard procedures (Brenner, 1974) and are listed in Table S1.

### Sample preparation

For the analysis of L1 larvae, gravid adults were bleached in bleaching solution (71% 1 M NaOH, 29% NaOCl), washed 4 times with M9, and the embryos allowed to hatch overnight in M9. L1 larvae were anesthetized using 100 mM NaN_3_ in M9, placed on a 2% agarose pad, covered with an 18 x 18 mm coverslip sealed with VaLaP (1:1:1 mixture of petroleum:jelly:lanolin:paraffin wax) and imaged within 1 hour. To obtain germ cell-free adults, *glp-4(bn2ts)* embryos were allowed to hatch at 16°C for one day and subsequently shifted to 25°C. The resulting adult worms were ethanol-fixed as described below.

### Microscopy and analysis

Widefield microscopy was performed on a Zeiss Axio Observer D.1 inverted microscope equipped with a 63 x Plan-Apochromat (NA 1.4) objective, connected to an Andor Zyla 4.2 sCMOS camera and an LED light source (Lumencor SOLA II) and controlled by the open-source µManager software (Edelstein et al., 2014). Pixels were binned 2x2. Confocal images were acquired on an upright Leica SP8 with 2 hybrid photon counting detectors (HyD) and a transmission photomultiplier tube (PMT) for brightfield; imaging was carried out with a 63 x HC Plan-Apochromat (NA 1.4), using 405, 488 and a 552 nm solid-state laser light for excitation and a DFC 7000 GT (B/W) camera. The confocal microscope was set to 8 kHz resonance scanning mode, 8 x line averaging and 2 x frame accumulation with a pixel size of 81 nm and the pinhole was set to 1 Airy units. Notch filters were used.

Single time point images of L1 larvae were acquired in 0.5 µm steps. For the analysis of adults, focal planes were acquired in 0.7 µm Z steps. Images were rotated, z-projected and brightness/contrast adjusted for each channel using Fiji (ImageJ), keeping the same settings within a series unless indicated otherwise. Adult images were stitched together in a pairwise fashion before identification of cells.

### Identification of cells in the L1 larva and adult

The number of foci was determined manually, counting twice, with an average difference between repeats of 11 % (+/-7 StDev) for GFP::SAS-7 and 8 % (+/-7 StDev) for SAS-4::GFP (see Fig. 2B for average foci number). For SAS-4::GFP, one outlier was removed in which seam cells in the head had already proliferated. For SPD-5::GFP, two outliers were removed due to a large excess of additional foci (in these outliers the numbers could not be counted manually but likely exceeded 200 foci compared to 116 on average for the worms included in the dataset. Moreover, these foci also formed larger blobs, for reasons that are unclear.

**Figure 2:**
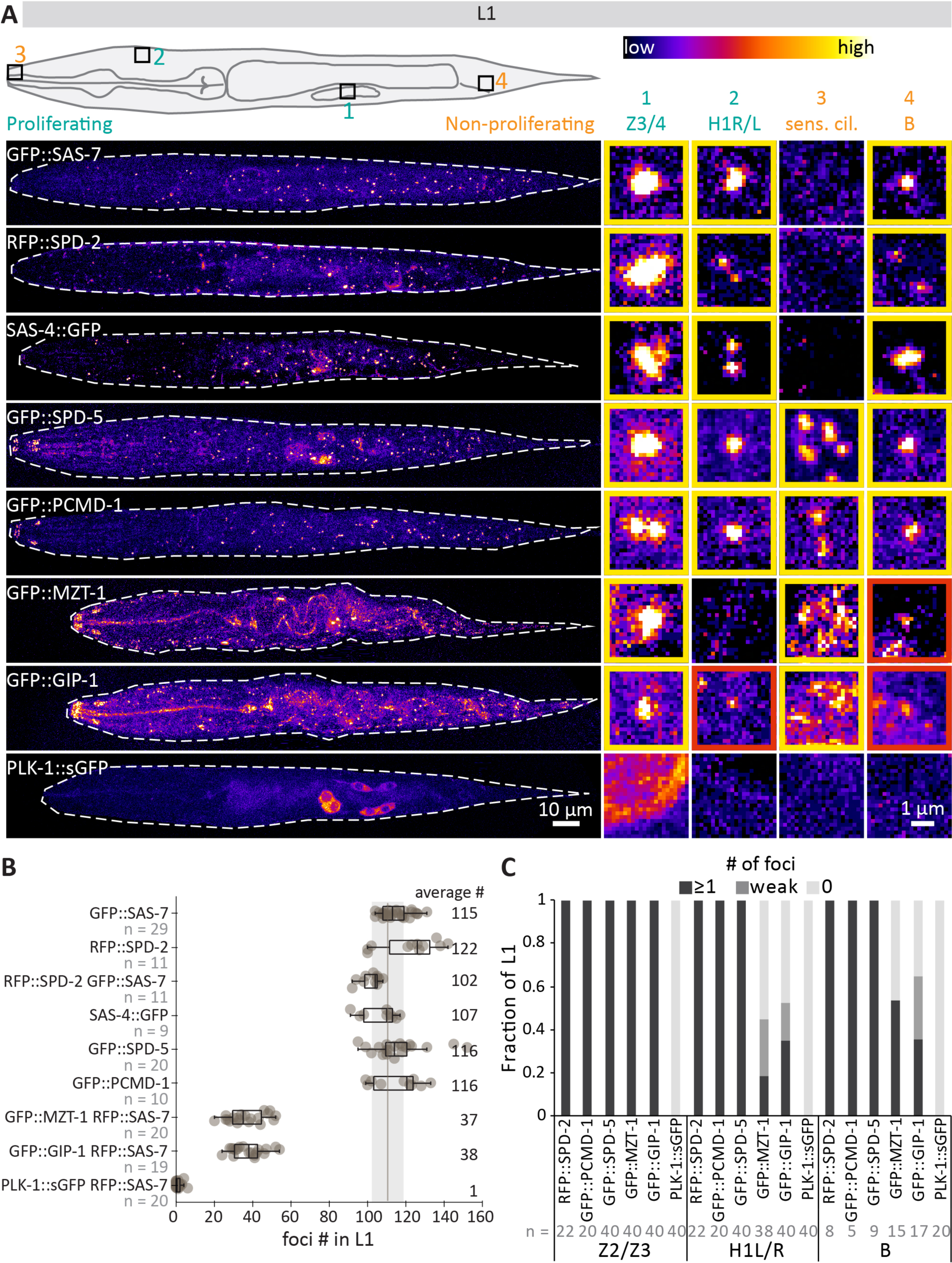
Some PCM core components are present at centrioles in L1 larvae. **(A)** Schematic of L1 larva with cells of interest highlighted by numbered squares, as well as maximum z-projections confocal microscopy of live paralyzed L1 larvae expressing the indicated markers. Proliferating cells are marked in turquoise, non-proliferating cells in orange. Insets show select planes containing centrioles in cells of interest highlighted in the schematic. Note that for GFP::MZT-1, GFP::GIP-1 and PLK-1::sGFP, the position of the centrioles was guided by the RFP::SAS-7 signal. Note also that the RFP::SPD-2 signal is considered to reflect its presence at centrioles in most cells, not at the PCM, as the signal is generally more restricted than it is in Z3/Z4, which are in the G2 phase of the cell cycle and thus harbor a maturing PCM. Yellow contours indicate clear foci, red contours weak foci. **(B)** Number of foci detected with the indicated markers in individual L1 larvae. Numbers on the right indicate the average. Note that head sensory cilia, which are positive for GFP::PCMD-1, GFP::SPD-5, GFP::MZT-1 and GFP::GIP-1, are not included in the reported counts given the high density of these foci in head sensory cilia, corresponding to acentriolar centrosomes. **(C)** Fraction of non-proliferating cells maintaining indicated centriolar markers. The number of cells analyzed is indicated in grey. Note that SAS-4::GFP and GFP::SAS-7 are always present in these cells (Kalbfuss and Gönczy, 2023).

Centrioles of non-proliferating cells were identified based on their stereotyped position, as the first centrioles visible on the left and right side in the head (AmsoR/L) (Kalbfuss and Gönczy, 2023), or around the rectal slit, which was visible in transmitted light. In the adult, tissues were identified based on their reproducible positions and nuclear characteristics (Kimble & Hirsh, 1979; Sulston & Horvitz, 1977). The identity of somatic gonad cells was verified based on the tissue-specific marker *pha-4p*::HIS-24::mCherry.

We noticed that using *glp-4(bn2ts)* in combination with SAS-4::GFP resulted in animals with mild phenotypic manifestations at 25°C, including a more bloated body size and defective intestinal nuclei. Therefore, we monitored SAS-4::GFP persistence at 20°C in worms of genotype *sas-4(bs195[sas-4::gfp] III, glo-1(zu931)X, bqSi189[pBN13(unc-119(+) Plmn-1::mCherry::his-58)] II*. In this background, whereas spermatheca and uterus cells can be readily identified, this is less clear for sheath cells, which were therefore not analyzed.

### Ethanol fixation

Adult worms were ethanol-fixed as described (Woglar et al., 2022), which resulted in a clear centriolar signal. Briefly, adult worms were washed off the plate into a reaction tube using PBS + 0.05% Tween-20 (PBS-T), and then washed once using PBS-T. After removing most of the PBS-T, 1 ml 100 % EtOH was added and the tube incubated for 3 min at room temperature. After removing all EtOH, 50 µl of 1:1 dilution M9 + Vectashield containing 1 µg/ml Hoechst 33258 was added. After rehydration for 1 min, worms were mounted on a slide with coverslip.

### Fusion experiments using AFF-1 or EFF-1

To induce fusion, L1 larvae of genotypes *hyEx173 [hsp16-2::aff-1 + ajm-1::GFP + rol-6(su1006)], glo-1(zu931)X, sas-7(is1[tagRFP::sas-7+loxP])III,* or *hyEx21 [hsp-16.2p::eff-1 + rol-6(su1006)], muIs65 [ajm-1::GFP + dpy-20(+)], glo-1(zu931)X, sas-7(is1[tagRFP::sas-7+loxP])III)*, or *sas-7(is1[tagRFP::sas-7+loxP])III, glo-1(zu931)X, wIs54 [scm::GFP] V, hyEx173 [hsp16-2::aff-1 + ajm-1::GFP + rol-6(su1006)])* were heat-shocked at 32°C for 30 min in a water-bath on plates with OP50, and subsequently incubated for the indicated times.

### Lineage tree analysis

For cumulative age of tissues, the time of the last cell division before differentiation was determined based on the lineage trees of the hermaphrodite and of the male (Kimble & Hirsh, 1979). Subsequently, cumulative births plots were generated using the python lifelines package (Davidson-Pilon, 2019).

### Statistics

Statistics were performed in R and are indicated in the figure legends. For significance tests, a Shapiro test for normality was performed. If no normality was found, a Wilcoxon-Mann-Whitney test was performed and Bonferroni corrected if multiple distributions were compared. Differences in variance were computed by the F-test, and if the F-test showed differences, a Welch test was performed. Otherwise, a standard Student’s t-test was performed. Box plots show median; top and bottom of the plots are the 75^th^ and 25^th^ percentile of the data. Whiskers extend to the maximum/minimum set of data sets that is less than 1.5 times the interquartile range. Everything beyond is considered an outlier and usually plotted unless otherwise indicated.

## Results

### The centriole elimination program is not constantly active in L1 larvae

The stereotyped fate of centrioles in L1 larvae, with some cells always retaining foci of centriolar proteins and other cells always losing them (Kalbfuss and Gönczy, 2023), prompted us to address whether there is an activity present in the latter set that is sufficient to eliminate centrioles. If this were the case and if this activity was diffusible, then fusing cells that normally retain centrioles with cells that normally lack them should result in organelle removal from the former set (Fig. 1B, left). Alternatively, in the absence of such an elimination activity, cells that normally retain centrioles should do so even following fusion (Fig. 1B, right).

To experimentally provoke cell fusion in L1 larvae, we induced the expression of the fusogens EFF-1 or AFF-1 using a heat shock promoter (del Campo et al., 2005; Sapir et al., 2007), in animals that also express the centriolar marker GFP::SAS-7. The effectiveness of fusion was verified by loss of the epithelial junction marker AJM-1::GFP from seam cell plasma membranes (Fig. 1D, compare with Fig. 1C). Importantly, we found that despite extensive cell fusion between cells that normally retain centrioles and cells that normally do not, the number of RFP::SAS-7 foci in the mid-body region did not change 4, 8 or 24 hours after induction of fusion by heat-shock (Fig. 1E). These results indicate that there is no diffusible activity sufficient to eliminate centrioles in L1 larvae, suggesting that maintenance of centriole is the default fate at this stage of development.

### All centrioles in L1 larvae maintain the PCM core components PCMD-1 and SPD-5

Is the maintenance of centrioles marked by SAS-4::GFP and GFP::SAS-7 in 68 cells of L1 larvae accompanied by persistence of PCM core proteins? Note that either 1 or 2 foci of SAS-4::GFP and GFP::SAS-7 can be detected in each of these 68 cells, depending on whether the minute centrioles are sufficiently distant from one another to be distinguished as single entities (Kalbfuss and Gönczy, 2023). As a result, the 68 cells in question exhibit on average 108 (+/- 2 SD) (n=4 animals) foci of SAS-4::GFP, and 108 (+/-5) (n=7) foci of GFP::SAS-7 (Fig. 2B). As shown in Figure 2A and 2B, we found a very similar number when examining the dual centriolar and PCM core protein RFP::SPD-2, as well as the PCM core components GFP::PCMD-1 and SPD-5::GFP, excluding from this analysis foci in sensory cilia since these are known to be acentriolar centrosomes (Garbrecht et al., 2021; Magescas et al., 2021; Serwas et al., 2017). By contrast, the number of foci for the more peripheral PCM components GFP::MZT-1 and GFP::GIP-1 was only 37 (+/- 9) and 38 (+/- 8), respectively (Fig. 2A, 2B).

We likewise analyzed the distribution of the Polo-like kinase PLK-1. In *Drosophila*, the departure of Polo from centrosomes during oogenesis results in centriole elimination (Pimenta-Marques et al., 2016). Therefore, we wondered whether in *C. elegans* the persistence of centrioles in 7 non-proliferating cells in L1 larvae could reflect the presence of PLK-1. However, foci of endogenously tagged PLK-1::sGFP could essentially not be detected anywhere at this stage of development (Fig. 2A-C, S1A, S1B), including in these 7 cells, although the fusion protein localizes to centrioles during embryogenesis (Martino et al., 2017). In L1 larvae, a strong PLK-1::sGFP signal was detected systematically in the cytoplasm of the germline precursor cells Z2/Z3, with a weaker signal also spotted sometimes in a couple of neighboring cells (Fig. 2A). Overall, based on these distributions, we conclude that centriolar PLK-1 is unlikely to be required for the maintenance of centrioles in either non-proliferating or proliferating cells of *C. elegans* L1 larvae.

Why is the number of foci bearing GFP::MZT-1 and GFP::GIP-1 reduced compared to that harboring SAS-4::GFP and GFP::SAS-7 in L1 larvae? We wondered whether this reduction might reflect the fact that some cells are in the early stages of the cell cycle, when centrosomal GFP::GIP-1 and GFP::MZT-1 are expected to be weak or absent. By contrast, SAS-4::GFP and GFP::SAS-7 persist at centrioles throughout the cell cycle, so that their signal intensities should hardly vary. Compatible with this possibility, we found that H1L/R cells, which generally do not exhibit condensed DNA and thus are generally not nearing mitosis, usually do not harbor a focus of GFP::GIP-1 or GFP::MZT-1 (Fig. 2A, 2C). By contrast, the germ line precursor cells Z2/Z3, which exhibit condensed DNA and are in the G2 phase of the cell cycle (Fukuyama et al., 2006), harbor GFP::MZT-1 and GFP::GIP-1 foci (Fig. 2A, 2C). Overall, our observations suggest that only a subset of centrosomes exhibit MTOC activity in L1 larvae, probably as a function of cell cycle state, as exemplified by the comparison of GFP::GIP-1 and GFP::MZT-1 distribution between H1L/R and Z2/Z3.

Next, we examined specifically the status of PCM proteins in the subset of non-proliferating cells that maintain SAS-4::GFP and GFP::SAS-7 foci in L1 larvae, except for the K’ cell, which is difficult to identify with certainty without dedicated nuclear markers. This analysis established that all 6 non-proliferating cells analyzed maintained the PCM core components RFP::SPD-2, GFP::PCMD-1 and SPD-5::GFP (Fig. 2A, 2C; Fig. S1A, S1B). By contrast, the outer PCM components GFP::GIP-1 and GFP::MZT-1 were generally not present in a focus, as anticipated from these 6 cells having terminally exited the cell cycle (Fig. 2A, 2C; Fig. S1A, S1B).

Overall, we conclude that centrosomes in non-proliferating cells and in proliferating cells that are paused in the G1 phase of the cell cycle harbor similar centriolar and PCM core proteins.

### Centriolar protein foci are present in specific terminally differentiated cells in the adult hermaphrodite

Not only is it the case that the majority of cells in L1 larvae that maintain centrioles proliferate later in development, but in addition 6 of the 7 non-proliferating cells with centrioles in the hermaphrodite are cells whose relatives are blast cells in the male (Kalbfuss and Gönczy, 2023). Therefore, it is possible that only those cells that will later proliferate in the hermaphrodite, or whose relatives in males will do so, retain centrioles. To explore whether other cells that have terminally exited the cell cycle retain centrioles in *C. elegans*, we addressed whether this is the case in such cells in the adult hermaphrodite. We used *glp-4(bn2ts)* temperature-sensitive mutant animals for these experiments to deplete proliferating germ cell nuclei, which harbour centrioles, thereby enabling us to readily identify all somatic cells that might maintain centrioles. *glp-4(bn2ts)* mutant animals are healthy, with apparently normal somatic gonads (Beanan & Strome, 1992), and retain only very few germ cells that were marked in our experiments using *mex-5p*::GFP::HIS-58 (Fig. 3A, Fig. S2A). We monitored simultaneously the distribution of GFP::SAS-7 and RFP::SPD-2 as a proxy for centrioles in young adult hermaphrodites, and found foci of both fusion proteins to be present in all animals (Fig. 3A-F, Fig. S2A, S3, S4). Subsequently, we identified individual tissues and scored whether specific cells herein harbour foci of both GFP::SAS-7 and RFP::SPD-2. Importantly, whereas most somatic tissues only exhibited sporadic instances of GFP::SAS-7 and RFP::SPD-2 foci (Fig. S3), we found many such foci in a systematic manner in the region of the somatic gonad (Fig. 3A-F, Fig. S4). In particular, we found that spermatheca and spermathecal-uterine junction cells always harbour cells with GFP::SAS-7 and RFP::SPD-2 foci (Fig. 3D, S4; n=7 animals). By contrast, we found that only some of the cells in the sheath (71+/-12% cells) and uterus (75 +/-15%) regions harbour colocalizing foci (Fig. 3F, Fig. S4). Nevertheless, cells in these two tissues generally maintain a GFP::SAS-7 focus in 90 (+/-19)% and 93(+/-7)% of cells, respectively, albeit not always with a colocalizing RFP::SPD-2 focus.

**Figure 3:**
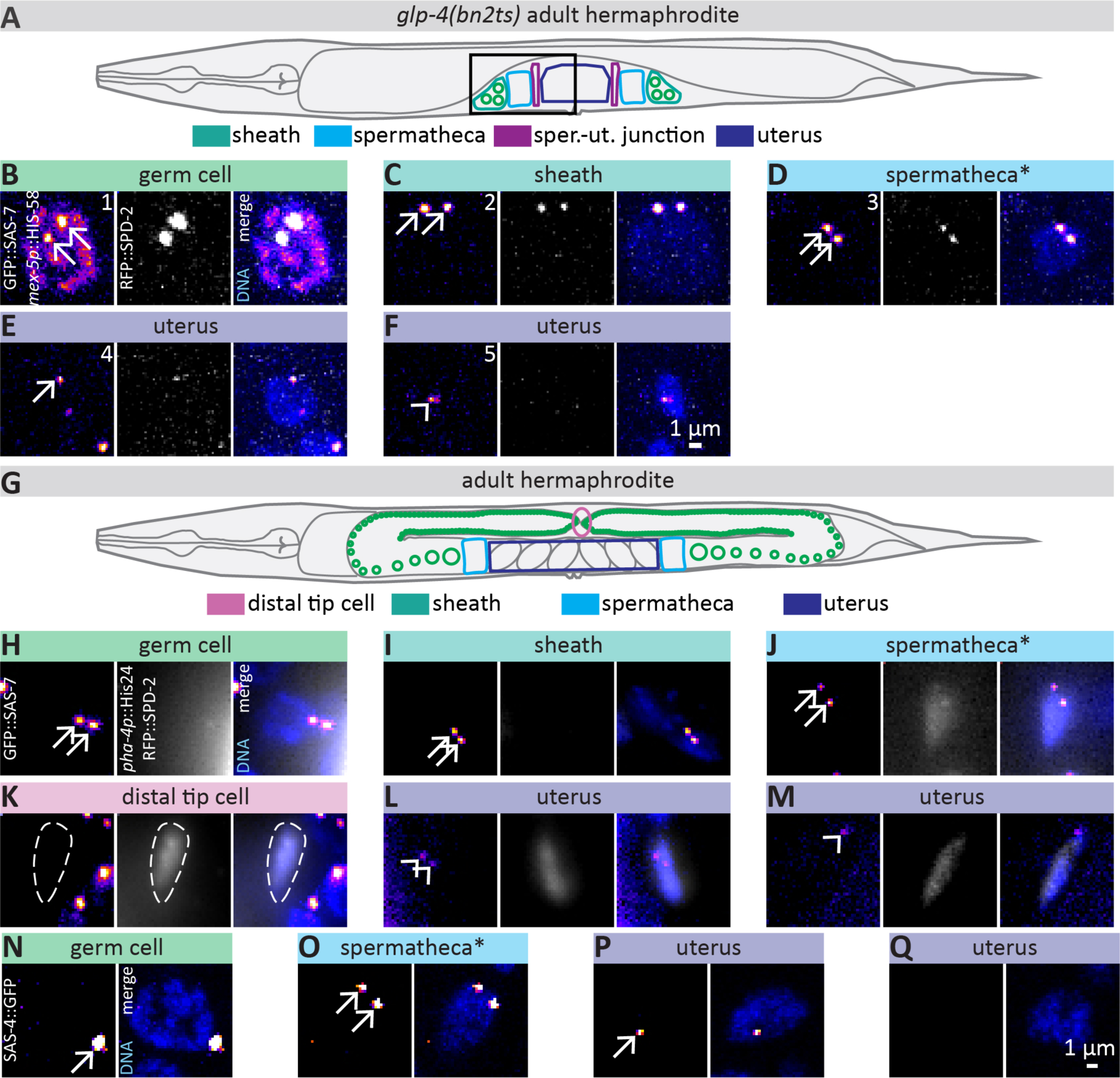
Centriolar foci are maintained in the somatic gonad of the adult hermaphrodite. **(A)** Schematic of *glp-4(bn2ts)* adult hermaphrodite grown at 25°C; black rectangle indicates with localization of somatic gonad organs. Green: germ line; turquoise: sheath cell; blue: spermatheca; violet: uterus. **(B-F)** High magnification of indicated cells of adult hermaphrodites imaged by confocal microscopy and maximum z-projected, showing relevant planes; see Fig. S2A for overview image. Note that most uterus cells maintain a weak focus of both GFP::SAS-7 and RFP::SPD-2 (E), and sometimes only a weak focus of GFP::SAS-7 (F), which was therefore not considered a proper centriole. Asterisk in (D) indicates that spermatheca and spermatheca-uterine junction cells could not be easily distinguished due to their proximity and are grouped here under the term “spermatheca”. **(G)** Schematic of wild-type adult hermaphrodite showing tissues of interest, the cells of which are highlighted in (H-Q). **(H-M)** High magnification images of indicated cells in *glp-4(bn2ts)* adult hermaphrodites grown at the permissive temperature, thereby yielding a worm without phenotype, expressing GFP::SAS-7, RFP::SPD-2, *pha4p*::His-24::mCherry, and stained with Hoechst to mark DNA. *pha4p*::His-24::mCherry was used as a somatic gonad marker, but was not detected in sheath cells, which were therefore identified based on position and morphology. *pha4* ::His-24 ::mCherry levels were adjusted for each cell. Arrows point to strong GFP::SAS-7 foci, arrowheads to weak GFP::SAS-7 foci. The distal tip cell nucleus is highlighted by a dashed contour in (K), and is in the vicinity of germ cells, with clear centriolar signal (right part of the panel). **(N-Q)** High magnification images of cells in adult hermaphrodites expressing SAS-4::GFP and stained with Hoechst. Tissues were identified based on their localization in the animal. Arrows point to SAS-4::GFP foci.

**Figure 4:**
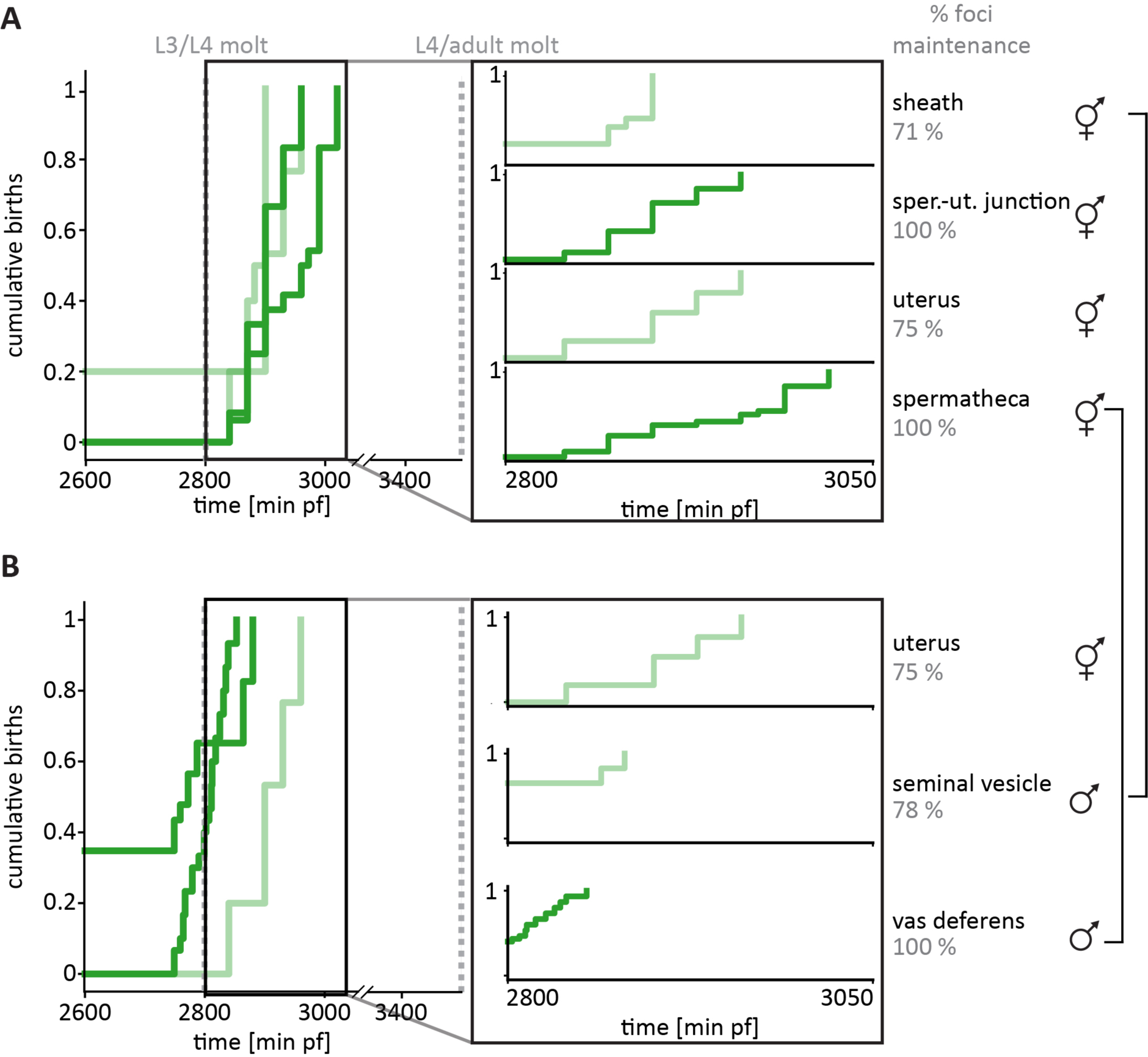
Centriolar foci are more reliably detected in tissues independent of their age. **(A-B)** Cumulative births of somatic gonad cells based on the lineage tree (Kimble & Hirsh, 1979); vertical black rectangle indicates time interval highlighted to the right for each individual tissue. Percentage of cells maintaining GFP::SAS-7 and RFP::SPD-2 are indicated. Dark green indicates that all cells in that adult tissue maintain GFP::SAS-7 and RFP::SPD-2 foci; light green indicates that this was the case in only some cells (see percentage to the right). (A) Among the hermaphrodite somatic gonad tissues, spermathecal cells are born last. Note that whereas spermatheca and spermatheca-uterine junction cells maintain GFP::SAS-7 and RFP::SPD-2 foci, the uterus and sheath cells harbor merely weak GFP::SAS-7 foci, and not in all cells (n=7 animals). (B) Although cells in the male vas deferens are born earlier than those in the uterus of the hermaphrodite, the former set retains foci of centriolar proteins whereas the latter set does not (n=3 animals). Note also that the recognition of cells belonging to the vas deferens is difficult in the absence of a tissue specific marker, such that the identification reported here should be considered merely as a best estimate. Functional homologs in hermaphrodites and males are indicated by brackets.

We set out to confirm these findings in worms with a normal germ line. To identify cells of the somatic gonad with increased certainty in this case, we used the somatic gonad marker *pha-4p*::His-58::mCherry (Kalb et al., 1998) (Fig. 3G-M). This enabled us to ascertain the presence of GFP::SAS-7 foci at spermatheca and spermatheca-uterine junction cells in 100% of cases, as well as in cells of the uterus in 95 +/-2% of cases (n=4 worms), in line with the above analysis. Using this marker, we found also that the distal tip cell does not maintain a focus of GFP::SAS-7 (Fig. 3K). We also assessed the presence of SAS-4::GFP in cells of the somatic gonad (Fig. 3 N-Q), finding SAS-4::GFP foci in 90 +/-5 % of cells in the spermatheca region (Fig. 3O), and in 48 +/-9% of cells in the uterus region (Fig. 3P, 3Q; n=5 worms). We note that these percentages are somewhat lower than those for the simultaneous presence of GFP::SAS-7 and RFP::SPD-2, raising the possibility that some foci may no longer be *bona fide* centrioles at the time of analysis. Regardless, the trend for foci maintenance per tissue type was the same for all markers.

Overall, we conclude that foci bearing signature centriolar proteins are maintained in some post-mitotic cells of adult hermaphrodites, most notably in the spermatheca of the somatic gonad.

### Centriolar foci are also maintained in the male somatic gonad

What could determine whether a post-mitotic cell of the adult hermaphrodite somatic gonad, derived from the Z1/Z4 precursor cells, maintains a centriole or not? We wondered whether this could reflect the age of the centriole, which conceivably could be eliminated at a set time after having been formed. To address this possibility, we plotted the cumulative time of birth of cells from the different tissue of the somatic gonad using the available hermaphrodite lineage tree (Kimble & Hirsh, 1979) (Fig. 4A). In these plots, a value of 1 means that all cells of the indicated tissue have been born at the indicated time. This analysis showed that the sheath is the tissue whose formation is completed first, whereas the spermatheca is completed last (Fig. 4A). Given that centriolar foci are systematically present in the spermatheca, this observation alone would be compatible with the notion that centriole age, which correlates with cell age, might determine centriole fate. However, time of birth analysis also showed that both spermatheca-uterine junction cells and the uterus are fully formed at similar times (Fig. 4A), even though only cells in the former systematically retain GFP::SAS-7 and RFP::SPD-2 foci, indicating a lack of strict correlation between centriolar age and fate.

To investigate further this question, we analyzed the presence of centriolar foci in the male. Like in the hermaphrodite, the Z1/Z4 lineages give rise to the complete somatic gonad in the male (Kimble & Hirsh, 1979). However, the timing and pattern of the lineage differs between the two sexes, both in terms of the tissues and the number of cells ultimately generated (Kimble & Hirsh, 1979). We determined and plotted the cumulative time of birth of cells from the male somatic gonad, finding that the male vas deferens, which is the functional homologue of the hermaphrodite spermatheca, and the male seminal vesicle, which is the functional homologue of the hermaphrodite sheath cells, are fully formed before the hermaphrodite uterus (Fig. 4B). Therefore, the male somatic gonad provides another opportunity to investigate whether centriole age might be key for organelle elimination. We found that foci harbouring both GFP::SAS-7 and RFP::SPD-2 foci persist also in adult males, being mostly restricted to the region encompassing the somatic gonad (Fig. 5A-E, S2B). Most salient was the fact that cells of the vas deferens invariably maintain GFP::SAS-7 and RFP::SPD-2 foci (Fig. 4B, 5D), despite them being born earlier than cells in the hermaphrodite uterus or in the male seminal vesicle, which sometimes do not harbour such foci (Fig. 4B). Interestingly, functional homologous tissues seem to have similar outcomes in terms of the percentage of cells maintaining foci per tissue type (compare Fig. 4A and Fig. 4B; sheath and seminal vesicle; spermatheca and vas deferens).

**Figure 5:**
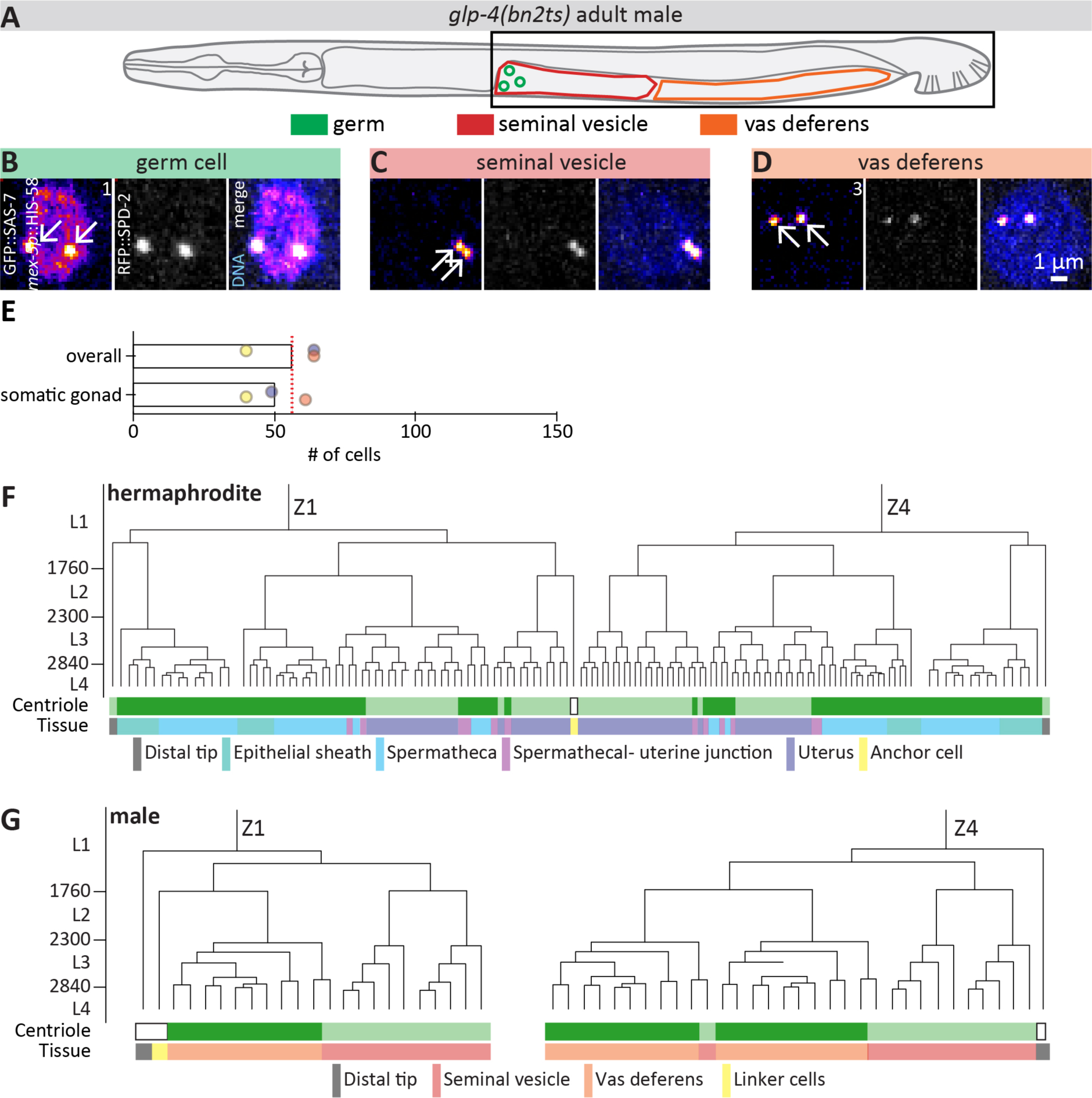
Centriolar foci are maintained in somatic gonad cells in the male. **(A)** Schematic of *glp-4(bn2ts)* adult male grown at 25°C, highlighting somatic gonad region where cells with colocalizing foci of GFP::SAS-7 and RFP::SPD-2 cells are present. The black rectangle in the schematic highlights the region shown in Fig. S2B. **(B-D)** Confocal microscopy maximum z-projections of cells in the region of germ cells (B), seminal vesicle (C) and vas deferens (D); only selected planes are projected. See Fig. S2B for overview image. **(E)** Number of cells in the male simultaneously maintaining foci of GFP::SAS-7 and RFP::SPD-2. Red dotted line indicates number of cells stemming from Z1/Z4 (56 cells). Note that the majority of cells maintaining centriolar foci in adults stem from the somatic gonad progenitor cells Z1/Z4. **(F, G)** Lineage tree of the somatic gonad in hermaphrodites (F) and males (G). Components always maintaining GFP::SAS-7 and RFP::SPD-2 foci are marked in dark green below the lineage tree, those sometimes without RFP::SPD-2 in light green. In addition, the type of tissues are indicated below the lineage tree.

Overall, our analysis in adult hermaphrodites and adult males reveals that centriolar foci are reliably detected in most somatic gonad cells (Fig. 5F, 5G), and indicates that their maintenance does not correlate with age of the cell, but rather with cell fate.

## Discussion

In this work, we show that a potential diffusible centriole elimination activity is not present in *C. elegans* L1 larvae. Furthermore, we find that all centrioles present at that stage maintain a PCM core consisting of PCMD-1 and SPD-5, but sometimes lack GIP-1 and MZT-1, potentially reflecting cell cycle differences. We also reveal that, in the adult, functionally equivalent terminally differentiated cells of the somatic hermaphrodite and male gonad retain centriolar foci, and that such retention does not simply reflect cell age, but instead cell type.

### L1 larvae do not contain a diffusible elimination activity sufficient to remove centrioles

Cell fusion experiments represent a powerful means to obtain insights into the mechanisms governing cellular processes, and have been instrumental for instance in revealing the logic of the eukaryotic cell cycle (Johnson & Rao, 1970; Rao & Johnson, 1970). Here, we have shown that centrioles normally destined to survive are not eliminated in the L1 larval stage following cell fusion with cells lacking centrioles. This observation indicates that L1 larvae do not contain a diffusible elimination activity sufficient to remove centrioles, although we cannot formally exclude that such an activity would be present, but short lived or poorly accessible by the cells containing centrioles. Overall, these observations lead us to propose that centriole elimination is an organelle-intrinsic process in this context. However, we note that the elimination of centrioles in cells that lack them in L1 larvae occurred already during embryogenesis, such that a temporally restricted cytoplasmic elimination activity would not have been revealed by fusion experiments conducted at the L1 stage.

Regardless, our observation of centriole elimination not being constantly active shows interesting parallels with a naturally occurring cell fusion: that of oocyte and sperm at fertilization. Centrioles are eliminated in diplotene of prophase I in *C. elegans* oocytes, (Mikeladze-Dvali et al., 2012), yet after fusion of oocyte and sperm a few hours thereafter, the two sperm-contributed centrioles are not eliminated in the newly fertilized zygote. Likewise, the last maternally-contributed centriole is not eliminated in starfish zygotes despite being in the same cytoplasmic milieu as the two paternally-contributed centrioles (Borrego-Pinto et al., 2016; Pierron et al., 2020). Furthermore, in physiological polyspermy, the oocyte fuses with multiple sperm cells, and except for one functional centrosome pair, the centrosomes of the other sperm cells are presumably inactivated and thus do not interfere with bipolar spindle formation (reviewed in Snook et al., 2011). Overall, these observations reinforce the notion that centriole-intrinsic mechanisms are at play, which might be “encoded” in the centriole before terminal differentiation, perhaps in the form of centriolar proteins and their post-translational modifications.

### PLK-1 localization indicates that it does not modulate post-embryonic centriole fate

During *Drosophila* oogenesis, loss of the Polo kinase leads to PCM loss and subsequent centriole elimination, whereas artificial tethering of Polo to the centrosome via the PACT domain prevents organelle demise (Pimenta-Marques et al., 2016). Here, we found that all centrioles detected in the L1 stage maintained PCM core components such as GFP::PCMD-1 and GFP::SPD-5. Surprisingly, however, PLK-1, the homologue of Polo, is absent from all centrioles in L1 larvae. These observations indicate that PLK-1 is not needed for centriole maintenance in this setting, although PCM components may nevertheless be important in a PLK-1-independent manner. Echoing these conclusions, experiments in the starfish *P. miniata* showed that pharmacological inhibition of Plk1 kinase does not lead to precocious centriole elimination (Pierron et al., 2020). Therefore, it appears that the mechanism observed in *Drosophila* oogenesis does not operate in a systematic manner throughout the animal kingdom to protect centrioles from elimination.

### GIP-1 and MZT-1 are not part of the PCM core in all tissues of *C. elegans*

Previously, the PCM core was defined as PCM components that are present around the centrioles also during interphase (Erpf et al., 2019). Analyses in the embryo (Magescas et al., 2019), the intestinal primordium (Sallee et al., 2018), where long gap phases are not yet present (Wong et al., 2022), and meiotic prophase germ cell nuclei (Woglar et al., 2022), where G1 cells are underrepresented (Fox et al., 2011), all suggest that GIP-1 and MZT-1 also localize to centrioles in interphase and therefore are part of the PCM core. Similarly, in human cells, the GIP-1 homologue GCP3 and the MZT-1 homologue MOZART1 localize to centrosomes throughout the cell-cycle (Hutchins et al., 2010; Murphy et al., 1998). By contrast, we could not detect these components on all centrioles in the L1 larval stage of *C. elegans*. Although we cannot exclude that the proteins are present below the detection limit, nor that analyzing foci in arrested L1 larvae might have helped reveal this feature given the longer time spent in a gap phase, together these observations suggest that GIP-1 and MZT-1 are not always part of the PCM core. In addition, since GIP-1 and MZT-1 are core components of the γ-TuRC, these findings raise the possibility that many centrosomes in the L1 larvae exhibit reduced or no MTOC activity.

### Centriolar foci maintained in adult tissues

We analysed the presence of foci bearing centriolar proteins in the adult hermaphrodite and the adult male, where all cells are post-mitotic, lest for those of the germline. Interestingly, we found that foci of centriolar markers are maintained also in specific adult tissues. In the absence of electron-microscopy data, and considering the variability of the signal intensity of the centriolar markers, we cannot unambiguously ascertain whether these foci correspond to *bona fide* centrioles or focused remnants of centriolar proteins. Regardless, interestingly, the maintenance of foci harbouring centriolar proteins was observed in functionally equivalent cells in hermaphrodites and males. Furthermore, the hermaphrodite uterus, where these foci were either absent or weak, is younger than all male somatic gonad tissues, indicating that centriole fate is not dependent on the age of the cell, but instead characteristic of the particular cell fate. This is in line with findings in the embryo, where centrioles are eliminated at different timepoints following mitosis in a cell fate-specific manner (Kalbfuss and Gönczy 2023). It remains to be understood why specific tissues maintain centriolar foci at least until a certain time in development, in contrast to the vast majority of terminally differentiated cells. One could speculate that this relates to their importance for signalling functions or cytoskeletal organization, which centrioles are known to orchestrate in other contexts (reviewed in Arquint et al., 2014; Muroyama & Lechler, 2017).

Overall, this work provides a map of centriole fate in the post-embryonic lineage of *C. elegans*, identifying cell-types that maintain foci of centriolar proteins past mitotic exit, thereby setting the stage for a comprehensive understanding of centriole fate and its regulation in the context of a developing organism.

## Acknowledgments

We thank Kevin O’Connell, Peter Askjaer, Jessica Feldman and Benjamin Podbilewicz for worm strains and insightful advice. We also thank Nikhil Bhatla for providing digital data of the hermaphrodite *C. elegans* lineage tree. Some strains were from the *Caenorhabditis* Genetics Center (CGC), which is funded by the NIH Office of Research Infrastructure Programs (P40 OD010440). We are grateful to Gabriela Garcia Rodriguez, Keshav Jha and Alexander Woglar for critical reading of a manuscript draft. Supported by a grant from the Swiss National Science Foundation (310030_197749 to PG).

## Author contributions

Conceptualization: NK, PG Methodology: NK, PG Investigation: NK, AB Writing: NK, PG Supervision: PG

## Funding

This work was supported by the Swiss National Science Foundation (grant 310030_197749 to PG). The funding source was not involved in study design, collection, analysis and interpretation of data, in the writing of the report and in the decision to submit the article for publication.

## Data availability

Data will be made publicly available on zenodo upon publication.

## Supplement

**Supplement 1:**
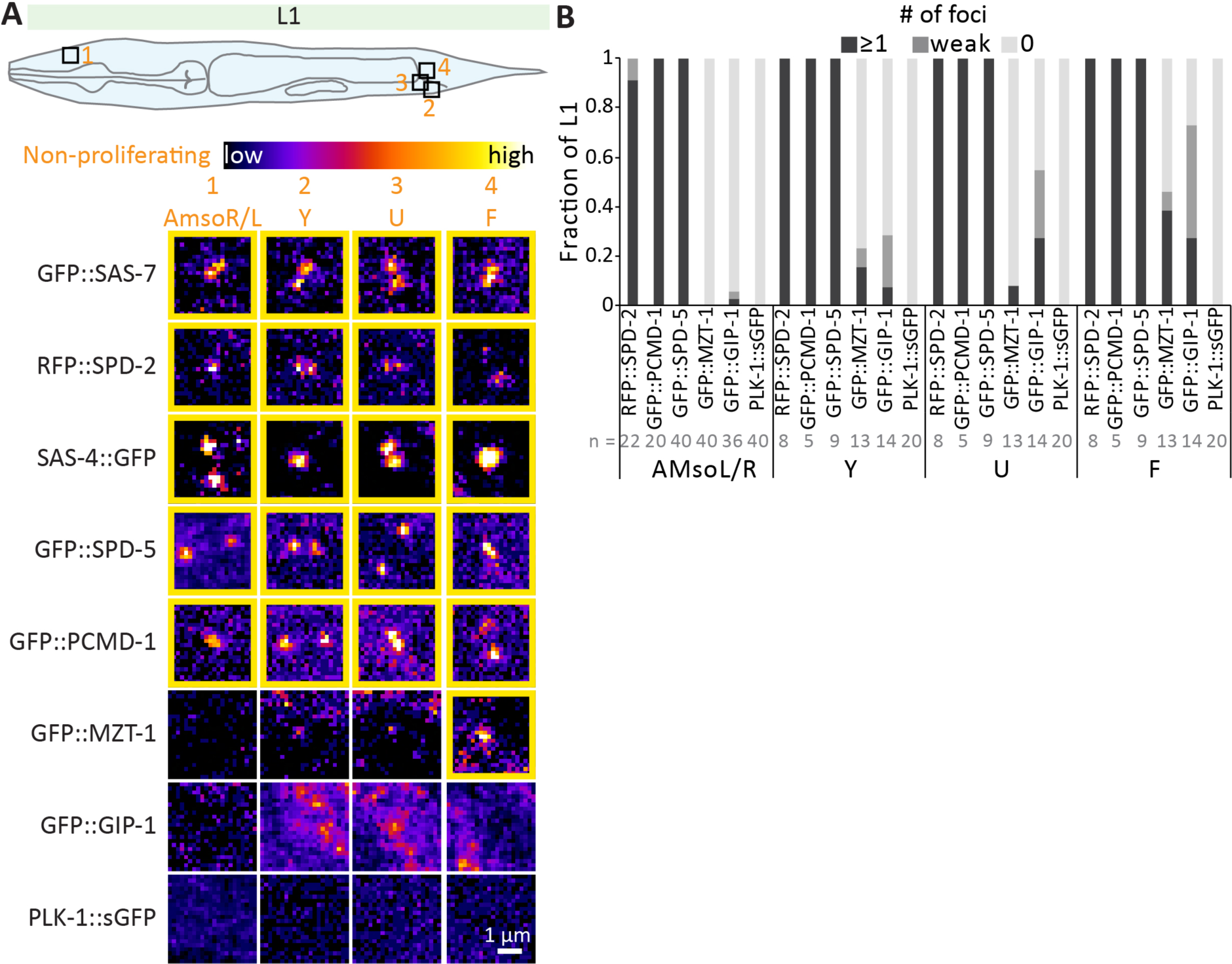
Non-proliferating cells maintaining centriolar foci also maintain some PCM core components. **(A)** Schematic of L1 larva with non-proliferating cells maintaining centrioles highlighted by numbered squares, as well as confocal microscopy maximum z-projections of live paralyzed L1 larvae expressing the indicated markers. Non-proliferating cells are marked in orange. Insets show select planes containing centrioles in cells of interest highlighted in the schematic. Note that for GFP::MZT-1, GFP::GIP-1 and PLK-1::sGFP, the position of the centrioles was guided by the RFP::SAS-7 signal. Yellow contours indicate clear foci, red contours weak foci. **(B)** Fraction of non-proliferating cells maintaining foci of indicated marker. Number of cells analysed indicated in grey. Note that SAS-4::GFP and GFP::SAS-7 are always present in all these cells (Kalbfuss and Gönczy, 2023).

**Supplement 2:**
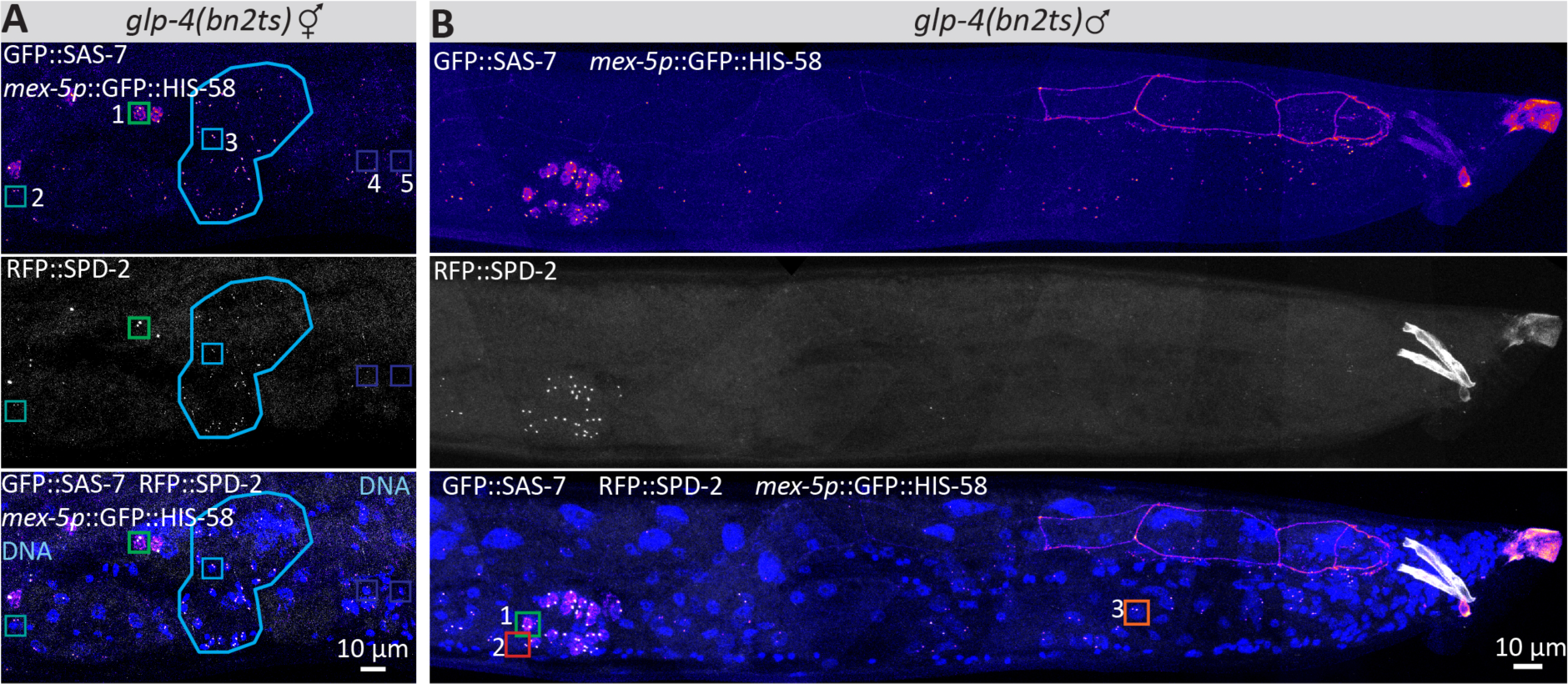
Overview of region with GFP::SAS-7 and RFP::SAS-7 colocalizing foci. **(A)** Confocal microscopy maximum z-projections of ethanol fixed adult hermaphrodite expressing GFP::SAS-7, RFP::SPD-2 and *mex-5p*::GFP::HIS-58 to identify the few remaining germ cells in *glp-4(bn2ts)* and stained with the DNA dye Hoechst. Colored numbered boxes indicate cells shown in Fig. 3B-F (green (1): germ cell; turquoise (2): sheath cell; blue (3): spermatheca; violet (4/5): uterus). The blue contour highlights the spermatheca, which serves as a convenient spatial landmark. **(B)** Confocal microscopy maximum z-projections of ethanol-fixed males stained with the DNA dye Hoechst and expressing GFP::SAS-7, RFP::SPD-2, and *mex-5p*::GFP::HIS-58 to identify the few remaining germ cells in *glp-4(bn2ts)*. Numbered and colored boxes indicate cells highlighted in Fig. 5B-D (green (1): germ cell; red (2): seminal vesicle; orange (3): vas deferens)

**Supplement 3:**
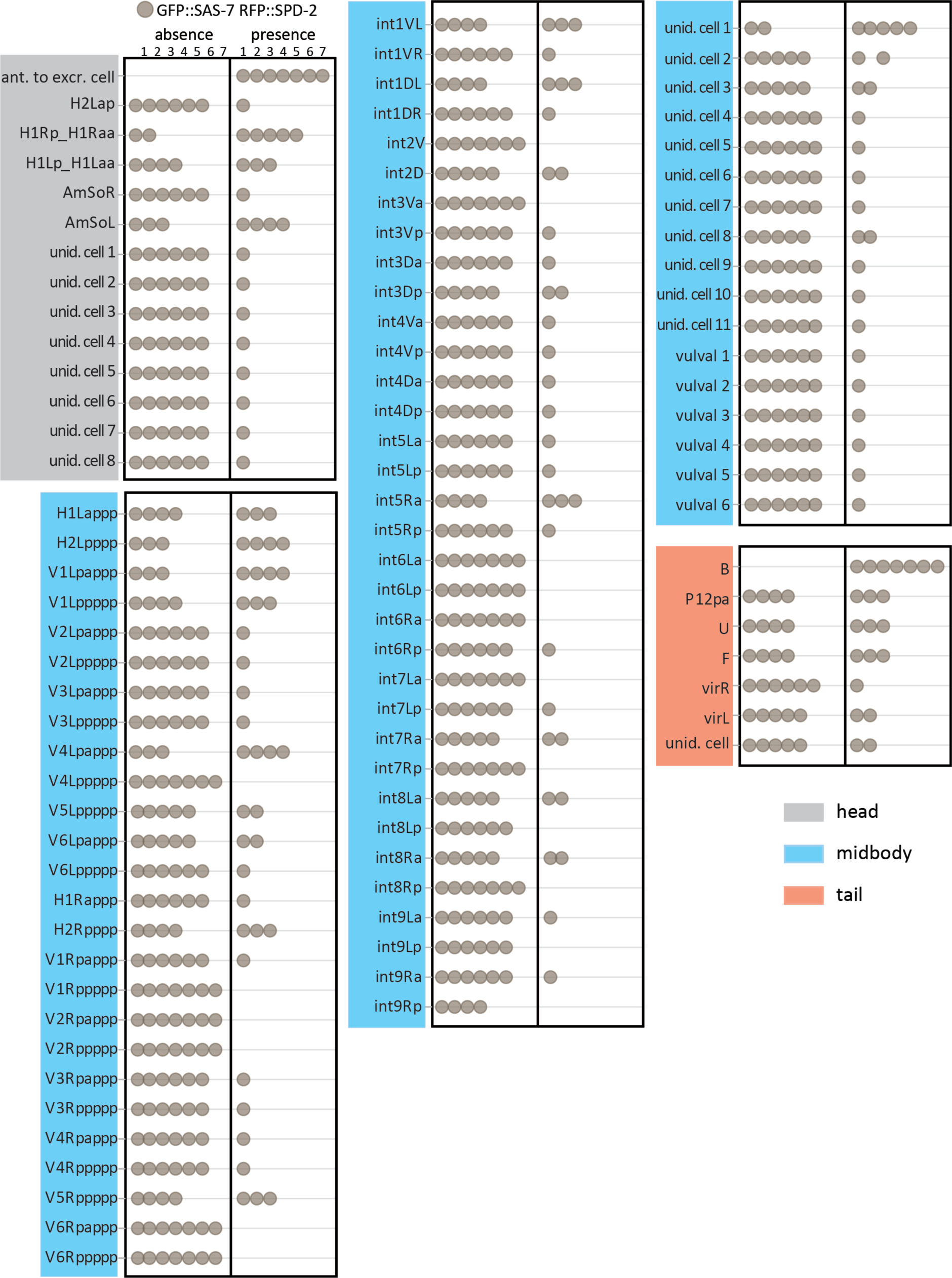
Somatic cells that maintain GFP::SAS-7 and RFP::SPD-2 foci sporadically. Worms expressing GFP::SAS-7 and RFP::SPD-2 (n=7), scored for the presence or absence of colocalizing foci in the indicated cells of the adult hermaphrodite. Note that in most of these cases, colocalizing foci were present only sporadically, suggesting that these are not *bona fide* centrioles. An exception is the B cell and a cell localized anteriorly to the excretory cell (ant. to excr. cell). Unid. Cell = unidentified cell. Regions of the hermaphrodite where foci were found are indicated in grey for head, blue for midbody and red for tail.

**Supplement 4:**
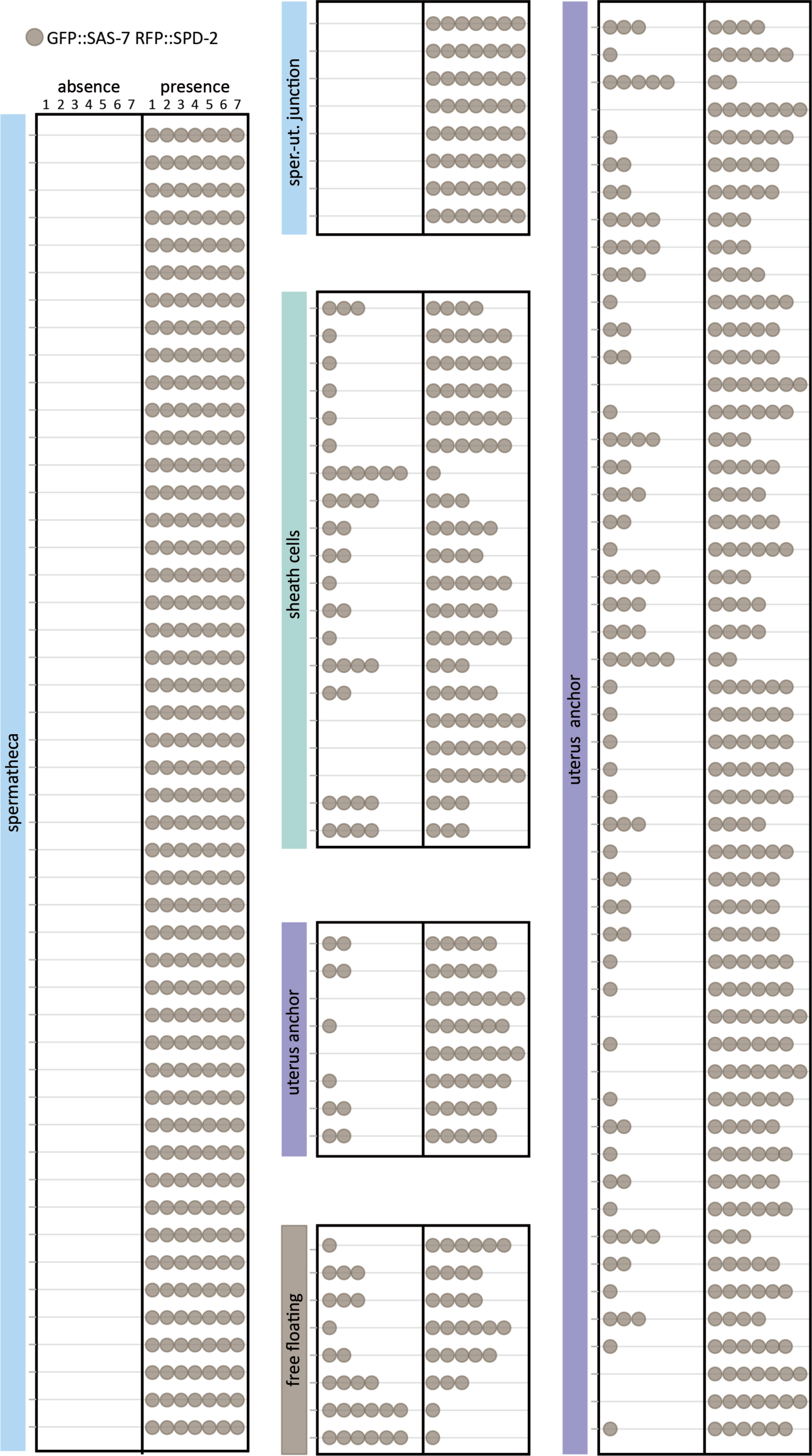
Stereotyped distribution of GFP::SAS-7 RFP::SPD-2 colocalizing foci in the spermatheca. Worms expressing GFP::SAS-7 and RFP::SPD-2 (n=7), scored for the presence or absence of foci in the indicated somatic gonad cells of the hermaphrodite adult. Tissue types were identified based on nuclear morphology and position. Individual cells were numbered going from anterior to posterior and dorsal to ventral. Cells between spermatheca and uterus were considered as cells of the spermathecal-uterine junction (sper.-ut. junction), although they could not clearly be distinguished from spermathecal cells, and are therefore represented jointly. Note that whereas the presence of these foci was stereotypical for spermatheca and spermatheca-uterine junction cells, cells of the other tissues sometimes did not harbor colocalising foci. When present, colocalizing foci in these tissues were also less bright than for spermatheca and spermatheca-uterine junction cells, suggesting that these might be centrioles in the process of being eliminated.

**Table S1:**
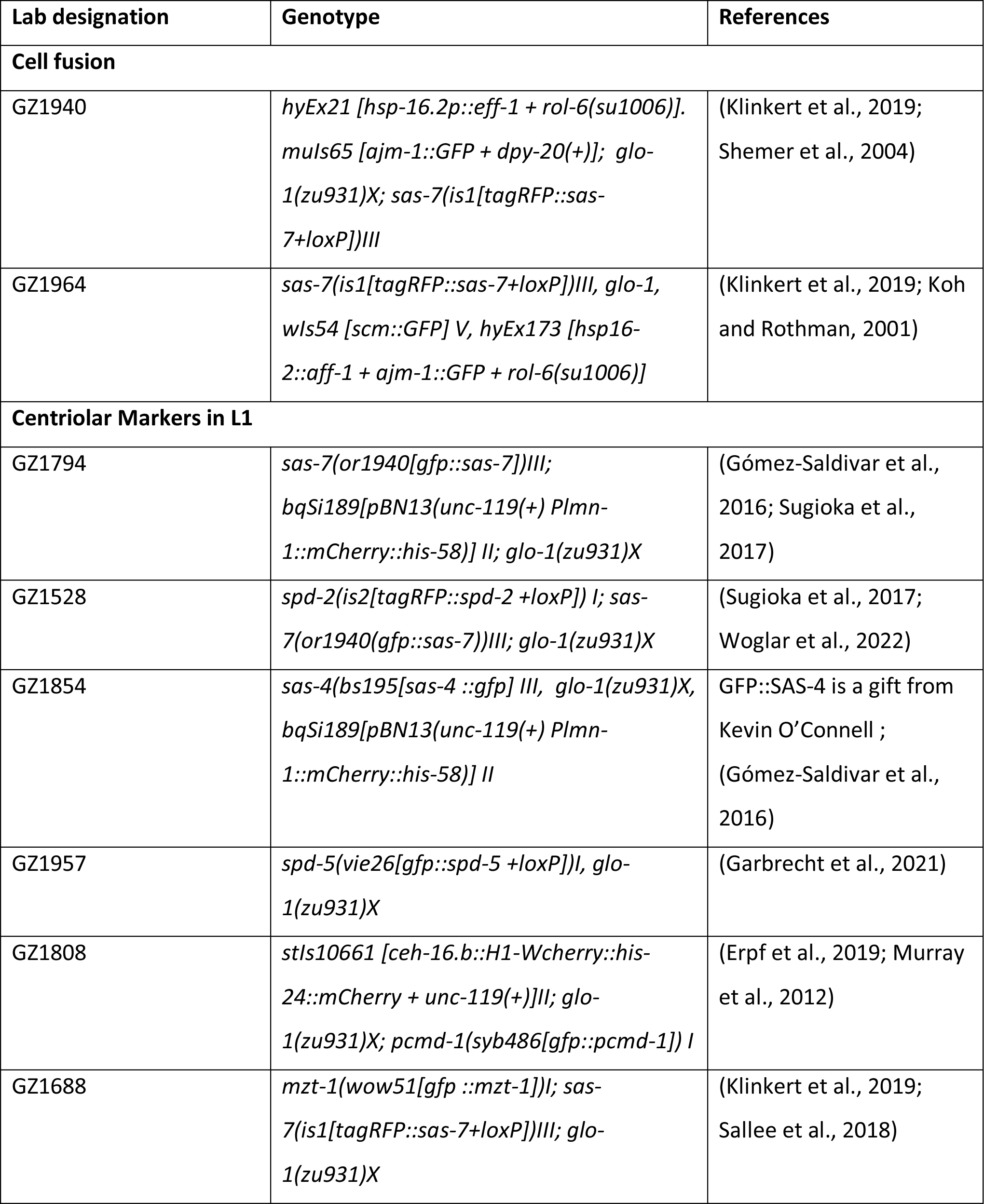
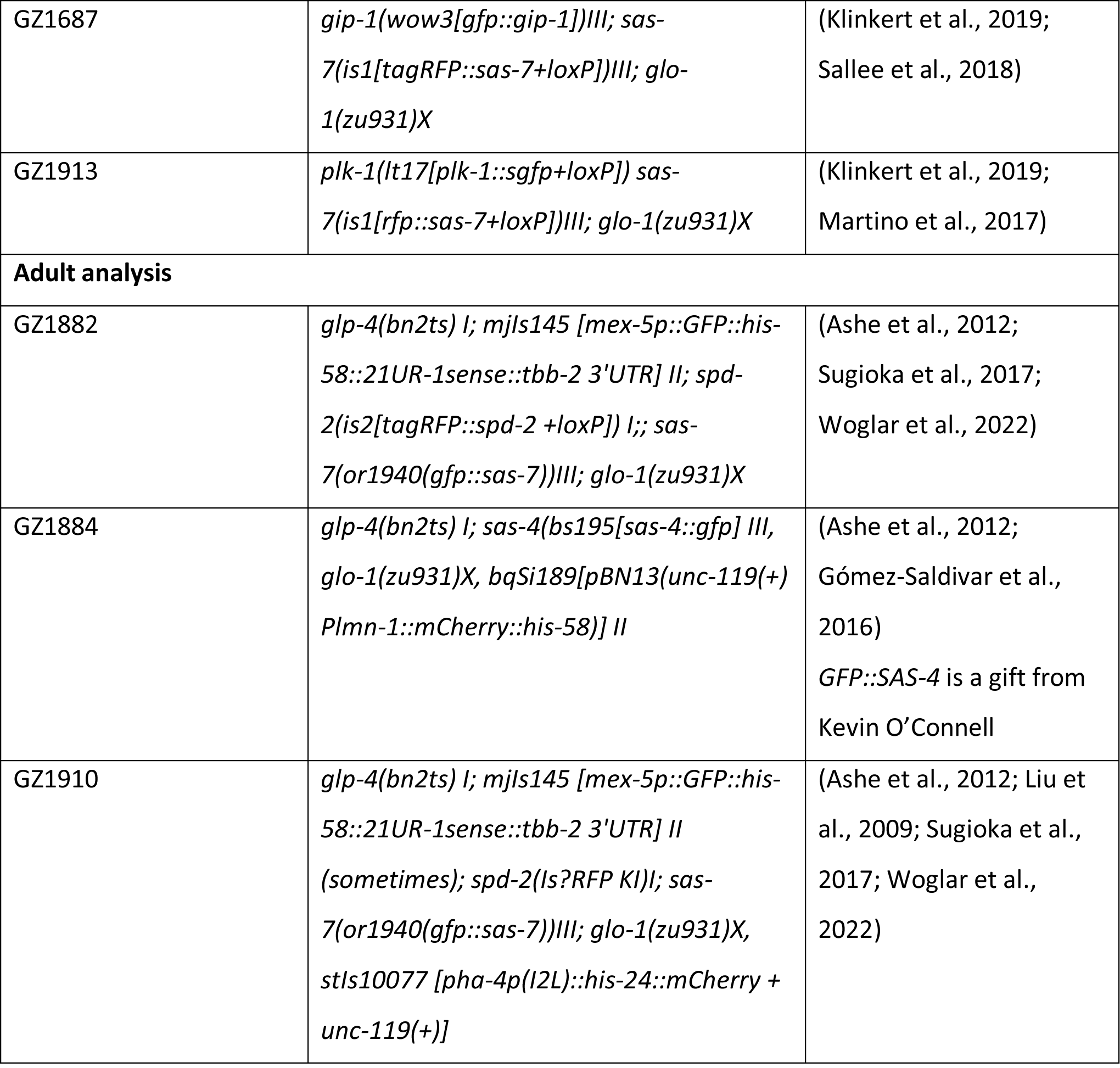
Strains used in this study.

## Notes

### Competing Interest Statement

The authors have declared no competing interest.

